# Dragotcytosis: Elucidation of the Mechanism for *Cryptococcus neoformans* Macrophage-to-Macrophage Transfer

**DOI:** 10.1101/387688

**Authors:** Quigly Dragotakes, Man Shun Fu, Arturo Casadevall

**Author notes:** Address Correspondence: Quigly Dragotakes Johns Hopkins Bloomberg School of Public Health Department of Molecular Microbiology and Immunology Baltimore, MD.

## Abstract

*Cryptococcus neoformans* is a pathogenic yeast capable of a unique and intriguing form of cell-to-cell transfer between macrophage cells. The mechanism for cell-to-cell transfer is not understood. Here we imaged macrophages with CellTracker Green CMFDA-labeled cytosol to ascertain whether cytosol was shared between donor and acceptor macrophages. Analysis of several transfer events detected no transfer of cytosol from donor to acceptor macrophages. However, blocking Fc and complement receptors resulted in a major diminution of cell-to-cell transfer events. The timing cell-to-cell transfer (11.17 min) closely approximated the sum of phagocytosis (4.18 min) and exocytosis (6.71 min) times. We propose that macrophage cell-to-cell transfer represents a non-lytic exocytosis event followed by phagocytosis into a macrophage that is in close proximity and name this process Dragotcytosis (Dragot is a Greek surname meaning ‘Sentinel’) as it represents sharing of a microbe between two sentinel cells of the innate immune system.

## Introduction

*Cryptococcus neoformans* is a pathogenic fungus that is the causative agent of cryptococcosis, a disease that affects primarily immunocompromised individuals. *C. neoformans* is a facultative intracellular pathogen that infects and reproduces inside of macrophages. Hence, the macrophage is a key cell in the pathogenesis of cryptococcosis and the outcome of the *C. neoformans*-macrophage interaction can determine the outcome of the infection(1–5). The cryptococcal pathogenic strategy is remarkable in that it involves fungal cell survival in a mature phagosome and the phenomenon of non-lytic exocytosis, which is characterized by expulsion of the fungal cells from the macrophage with the survival of both cells(6–8). In addition, *C. neoformans* is capable of being transferred from an infected to a non-infected macrophage(7, 8). Cell-to-cell transfer is generally believed to be a process different from non-lytic exocytosis, with these two events being referred to as Type III and Type II exocytosis, respectively(9), denoting the fact that all these events share in common the exit of a fungal cell from an infected macrophage. Non-lytic exocytosis has been described in mammalian(7, 8), fish(10), insect(11), and amoeba(12) cells and appears to be a highly conserved strategy for *C. neoformans* cells to escape host and environmental predatory phagocytic cells. Non-lytic exocytosis has been described in other pathogenic microbes, including *Burkholderia cenocepacia*(13), *Candida albicans*(14), and *Mycobacterium tuberculosis*(15), suggesting that it may be a widespread strategy for microbial escape from phagocytic cells.

Little is known about the mechanism of cell-to-cell transfer, which could facilitate the spread of infection in anatomical sites where macrophages are in close apposition to one another, such as cryptococcal granulomas and infected lymph nodes. Macrophage to endothelial cell transfer of *C. neoformans* was described in blood-brain barrier models(16). Whether cell-to-cell transfer favors control of infection, or promotes it, it is likely to depend on the circumstances of the host-microbe interaction. For example, transfer of a single fungal cell between two macrophages would appear to be a debit for the host, since *C. neoformans* residence in macrophages is associated with host cell damage(17) and thus could damage two host cells. Conversely, transfer of fungal cells from a macrophage infected with many yeasts could help in the control of infection since it would reduce the multiplicity of infection per cell.

In this study, we investigated the mechanism of macrophage-to-macrophage *C. neoformans.* The results are interpreted as consistent with an initial non-lytic exocytosis event followed by phagocytosis by a nearby macrophage, thus implicating non-lytic exocytosis in cell-to-cell transfer

## Materials and Methods

### C. neoformans Strain and Culture Conditions

Cryptococcal cultures were prepared by inoculating 10 mL Sabouraud Dextrose Broth [SAB; Becton-Dickenson, Franklin Lakes, NJ] media with a stab of frozen *C. neoformans* var. *grubii* serotype A strain H99 stock. Cultures were incubated at 30°C shaking at 150 rpm for 2 d before use in infections.

### Macrophage culture

Bone-marrow derived macrophages (BMDM) were generated from hind leg bones of 5-to 8-wk-old co-housed C57BL/6 female mice (Jackson Laboratories, Bar Harbor, ME) or Fc receptor knockout (Fcer1g) mice (Taconic model 583) of the same age. For the macrophage differentiation, cells were seeded in 100 mm tissue culture-treated cell culture dishes (Corning, Corning, NY) in Dulbecco’s Modified Eagle medium (DMEM; Corning) with 20 % L-929 cell-conditioned medium, 10 % FBS (Atlanta Biologicals, Flowery Branch, GA), 2mM Glutamax (Gibco, Gaithersburg MD), 1 % nonessential amino acid [Cellgro], 1 % HEPES buffer [Corning], 1 % penicillin-streptomycin [Corning] and 0.1 % 2-mercaptoethanol [Gibco] for 6 -7 d at 37°C with 9.5 % CO_2_. Fresh media in 3 ml were supplemented on day 3 and the medium were replaced on day 6. Differentiated BMDM were used for experiments within 5 days after completed differentiation.

Murine macrophage-like J774.16 cells were maintained in DMEM with 10 % NCTC109 medium [Gibco], 10 % FBS, 1 % nonessential amino acid, 1 % penicillin-streptomycin at 37°C with 9.5% CO_2_. All murine work was done using protocols reviewed and approved by IACUC. All experimental work in this study was done with BMDM except for the high-resolution movie shown in Figure S1, which was filmed using J774.16 cells.

### Acquisition of Supplemental Video

J774.16 cells were seeded (5 × 10^4^ cells/well) on poly-D-lysine coated coverslip bottom MatTek petri dishes with 14mm microwell [MatTek Brand Corporation] in medium containing 0.5 µg/ml lipopolysaccharide [LPS; Sigma], 0.02 µg/mL (100 U/ml) gamma interferon [IFN-γ; Roche]. Cells were then incubated at 37 °C with 9.5 % CO_2_ overnight. On the following day, macrophages were infected with cryptococcal cells s (1.5 × 10^5^ cells/well) in the presence of 10 µg/ml monoclonal antibody (mAb) 18b7. After 2 h incubation to allow phagocytosis, culture was washed five times with fresh medium to remove extracellular cryptococcal cells. Images were taken every 4 min for 24 h using a Carl Zeiss LSM 780 confocal microscope with a 40 × 1.4 NA Plan Apochromat oil-immersion DIC objective and a spectral GaAsP detector in an enclosed chamber under conditions of 5.0 % CO_2_ and 37 °C. Acquisition parameters, shutters and focus were controlled by Zen black software [Carl Zeiss].

### Macrophage Infections and Videos for Cytosol Exchange and Inhibitor Trials

BMDMs were seeded (2.5 × 10^4^ cells/well) in MatTek dishes and 24-well plates [Corning]. Cells in one MatTek dish and one well from the 24-well plate were activated overnight (16h) with IFNy (0.02 µg/mL) and (0.5 µg) LPS. Cells in the MatTek dish were stained with CellTracker Green CMFDA [ThermoFisher] according to manufacturer’s protocol before infecting with Uvitex 2B (5 µm/mL; Polysciences Inc., Warrington, PA) labeled and 18B7 (Generated in lab; 10 µg/mL) or guinea pig complement (20%; MilliPore) opsonized *Cryptococcus* at an MOI of 1 for 1 h. Cells from the 24-well plate were labeled with CellMask Orange [ThermoFisher] according to manufacturer’s protocol, then raised and seeded over the MatTek dish after washing with 2mL fresh cell media three times to remove extracellular cryptococcal cells. Cells were incubated for 30 additional minutes to allow for adhesion before supplementing the plate with 2 mL fresh cell media. MatTek dishes were then placed under a Zeiss axiovert 200M 10X magnification, incubated at 37°C and 9.5% CO_2_, and imaged every 2 min for a 24 h period. Images were then manually analyzed to identify clear transfer events. For each transfer event donor and acceptor cells were outlined according to cell membranes visible in phase contrast. CellTracker and Uvitex 2B channel intensities were then collected for each pixel within the cell outline and compared using unpaired two-tailed t tests between pre and post transfer quantifications.

Receptor inhibitor experiments used a single MatTek dish with 5 × 10^4^ macrophages activated overnight with IFNy (0.02 µg/mL) and LPS (0.5 µg/mL). Cells were infected with *C. neoformans* at an MOI of 1 for 1 h followed by three washes of 2 mL fresh media to remove extracellular cryptococcal cells. Cells were incubated for 1h with aCD16/32 anti-Fc receptor antibodies (0.5 µg/mL; BD Biosciences 553142) and CD11b anti-complement antibodies (0.5 µg/mL; BD Biosciences 553308) or Cytochalasin D (1 µg/mL; Sigma) before beginning 24 h imaging under the Zeiss axiovert 200M 10X magnification at 37 °C with 9.5 % CO_2_ overnight to analyze total cellular exit events.

Fc receptor knockout experiments used a single MatTek dish with 5 × 10^4^ macrophages activated overnight with IFNy (0.02 µg/mL) and LPS (0.5 µg/mL). Cells were infected with *C. neoformans* at an MOI of 1 for 1 h followed by three washes of 2 mL fresh media to remove extracellular cryptococcal cells. Cells were then imaged for 24 h under the Zeiss axiovert 200M 10X magnification at 37 °C with 9.5 % CO_2_ overnight to analyze total cellular exit events. For inhibitor trials, cells were incubated for 1h with CD11b anti-complement antibodies (0.5 µg/mL) before beginning 24 h imaging under the Zeiss axiovert 200M 10X magnification at 37 °C with 9.5 % CO_2_ overnight to analyze total cellular exit events.

For phagocytosis and non-lytic exocytosis timing experiments were set up as above except using an MOI of 1 for 2 h at 4°C, then MatTek dishes were placed on the microscope stage, and images were immediately taken every 1 min for 24 h immediately after adding mAb 18B7 (10 µg/ml) directly to the dish.

### Quantification of Temporal Kinetics

Each type of event (phagocytosis, non-lytic exocytosis, and cell-to-cell transfer) was given a start and end based on the movie frame when beginning and ending was observed. The total number of frames spanning from start to end were counted to estimate the duration of the event. The start of phagocytosis was defined as the first frame in which a cryptococcal cell was attached to a macrophage cell, no longer free moving through the media. The end of phagocytosis was defined as the first frame in which it is undeniably clear that the cryptococcal cell has been fully engulfed and is no longer touching the plasma membrane. The start of non-lytic exocytosis was defined as the frame immediately prior to a cryptococcal cell within a macrophage moving toward the plasma membrane. The end of non-lytic exocytosis was defined as the frame in which that cryptococcal cell is fully outside of its host macrophage and no longer in contact with the plasma membrane. The start of a cell-to-cell transfer event was defined in the same way as non-lytic exocytosis, that is the frame immediately prior to movement toward the plasma membrane. The end of a cell-to-cell transfer event was defined in the same way as phagocytosis, that is the frame in which the cryptococcal cell is fully engulfed by the acceptor macrophage and is no longer in contact with the plasma membrane.

### Statistics

Statistical differences between dye channels for both cytosolic (CellTracker Green) and cryptococcal (Uvitex 2B) were determined by two-tailed unpaired t-test between cells pre and post transfer. The region of interest was manually defined by the exterior of the host cell plasma membrane via phase contrast channel. Each pixel within the designated region was measured for intensity in its respective channel. Graphs are a depiction of pixel intensity values with bars representing minimum and maximum values. Statistics were calculated on these groups of pixel intensity values. For inhibitor trials significance was calculated using a one-sided test of proportions for each condition compared to the control (18B7 opsonized with no inhibiting antibody treatment).

## Results

### High Resolution Imaging of Cell-to-Cell Transfer

Cryptococcal cell-to-cell transfer was observed by imaging J774.16 cells infected with *C. neoformans* overnight with phase contrast microscopy (SMovie 1). The coordination apparently involved in this event suggests an underlying mechanism involving both the donor and acceptor cell.

### Cytosol transfer was not detected during cryptococcal transfer

To determine whether cytosol was transferred from donor to acceptor cell along with *Cryptococcus* cells during fungal macrophage-to-macrophage cell transfer, we visualized transfer events in which only the donor cell was stained with a permanent cytosolic dye in BMDMs. BMDMs were used in this and all proceeding experiments described in this manuscript. Upon identifying a transfer event the microscopic images were isolated before, during, and after the transfer event (Figure 1). This experiment was repeated until ten independent events were identified and the individual fluorescent channels were quantified at each frame before and after Cryptococcal cell transfer. We found that cytosolic dye signal remained constant in the donor (Figure 2A) and was not observable above background in the acceptor (Figure 2B) cell throughout the event. However, fluorescence intensity corresponding to cryptococcal cells decreased in the donor cell (Figure 2C) and increased in the acceptor cell (Figure 2D) after transfer. This analysis was repeated for every event (SFig 1). Taken together these data showed no evidence that host cell cytosol was transferred during cryptococcal cell-to-cell transfer. Additionally, we performed experiments supplemented with a plasma membrane stain (CellMask Orange; ThermoFisher) and identified two transfer events. There was no intensity difference between plasma membrane staining before and after transfer, suggesting that there is no mixing between donor and acceptor cell plasma membranes during transfer events (SFig 2).

**Figure 1.**
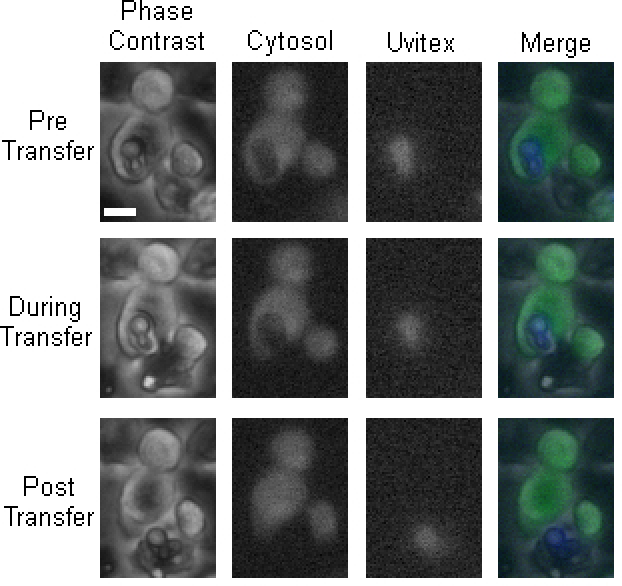
Dragotcytosis event captured via fluorescent microscopy. Representative frames before (Pre), during, and after (Post) cryptococcal transfer. 10X Phase Contrast, Cytosolic CellTracker Green CMFDA (Green), Uvitex (Blue), and merged channels. The Scale bar represents 10 µm and is constant for all images. Pre, During, and Post Transfer images were obtained from movie frames 266, 274, and 278, respectively. These images are from Event 4 in subsequent graphs.

**Figure 2.**
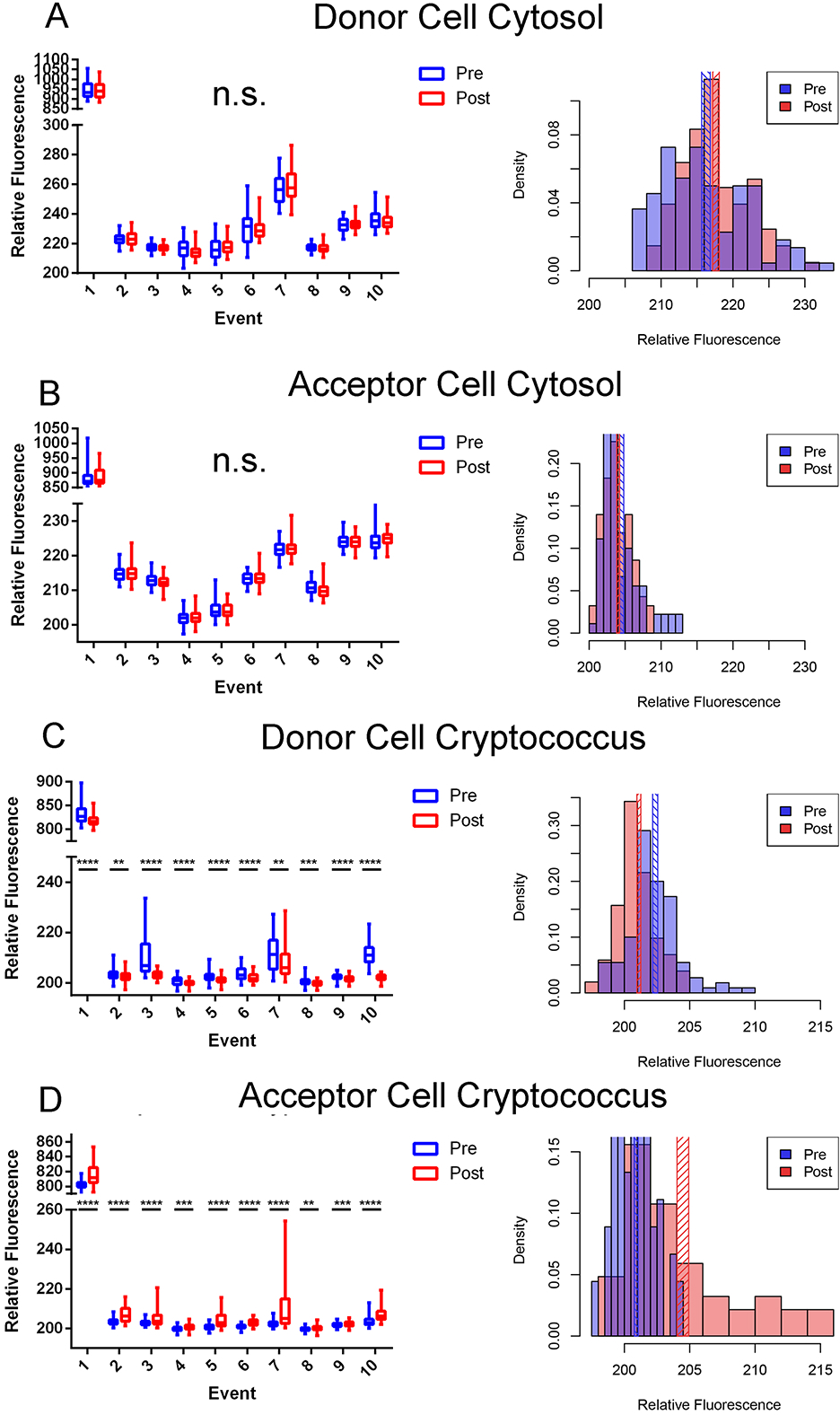
Quantifications of stains from Dragotcytosis events in biological replicates. **A.** Donor cell cytosol as measured by CellTracker Green CMFDA intensity before (blue) and after (red) transfer. **B.** Acceptor cell cytosol as measured by CellTracker Green CMFDA intensity before and after transfer. Intensity values remain at background levels in each replicate. **C.** Presence of cryptococcal cell inside Donor cell measured as Uvitex intensity before and after transfer. **D.** Presence of cryptococcal cell inside Acceptor cell measured as Uvitex intensity before and after transfer. Significance was determined by two tailed *t*-test with (****) representing a *P* value of < 0.0001, (***) representing a *P* value of <0.001, and (**) representing a *P* value of <0.01. Data for each population is each individual pixel intensity measurement, with bars representing minimum-maximum spans in box-whisker format. Density histograms represent population data specifically for Event 4 with cross-hatch rectangles representing mean +/-standard error. The relative fluorescence of Event 1 is higher than the other events because that measurement came from the initial experiment before dyes were titrated for optimization. Each of these events corresponds to a single event in which a single yeast is transferred from one macrophage to another with the exception of event 8 which spans two individual transfer events in which one yeast cell is transferred during each event. This was due to the necessity of acquiring clear frames for quantification.

### Cell-to-Cell Transfer Requires FcR and/or Complement Receptor

No cytosolic dye transfer between macrophages during transfer events suggests two hypotheses: 1. Cryptococcal cells are transferred via coordinated exocytosis followed by phagocytosis between the two macrophages; or 2. Only the phagosome is directly transferred between macrophages in a manner that excludes cytosol. Given that the *C. neoformans* capsule prevents phagocytosis(18, 19), that opsonin is required for ingestion(20), and that capsule-associated antibody is present after exocytosis(21), we designed experiments to test the first hypothesis. Specifically, we investigated whether cell-to-cell transfer was blocked by the addition of an anti-CD16/32 monoclonal antibody, which prevents FcR function, in BMDMs. Transfer events were drastically reduced by blocking the FcR but interestingly, occasional cell transfer events were still observed. It is known that opsonized cryptococcal cells can be phagocytosed via complement receptor (CR) by a mechanism where antibody modifies the capsule and allows direct interaction with this receptor in the absence of complement(22). To investigate whether this was the case, the experiments were repeated with complement inhibiting antibody (anti-CD11b) and both inhibiting antibodies. The frequency of cell-to-cell transfer was significantly reduced compared to control when each of the FcR and CR were inhibited and completely abrogated when both were inhibited together, with p < 0.01, 0.05, and 0.001, respectively (Figure 3A, Table I). To further explore this phenomenon, and whether the reduction in observed effects was an artifact of antibody incubation, we infected BMDMs harvested from Fc receptor knockout mice (Fcer1g) opsonized with 18B7 antibody. We found that Fcer1g BMDMs experienced cell-to-cell transfer significantly less than the control, p < 0.01, and at a similar rate as wild-type cells incubated with the anti-FcR antibody (Figure 3A, Table I). These data suggested that both Fc and complement receptor mediated phagocytosis can be utilized in cell-to-cell transfer.

**Figure 3.**
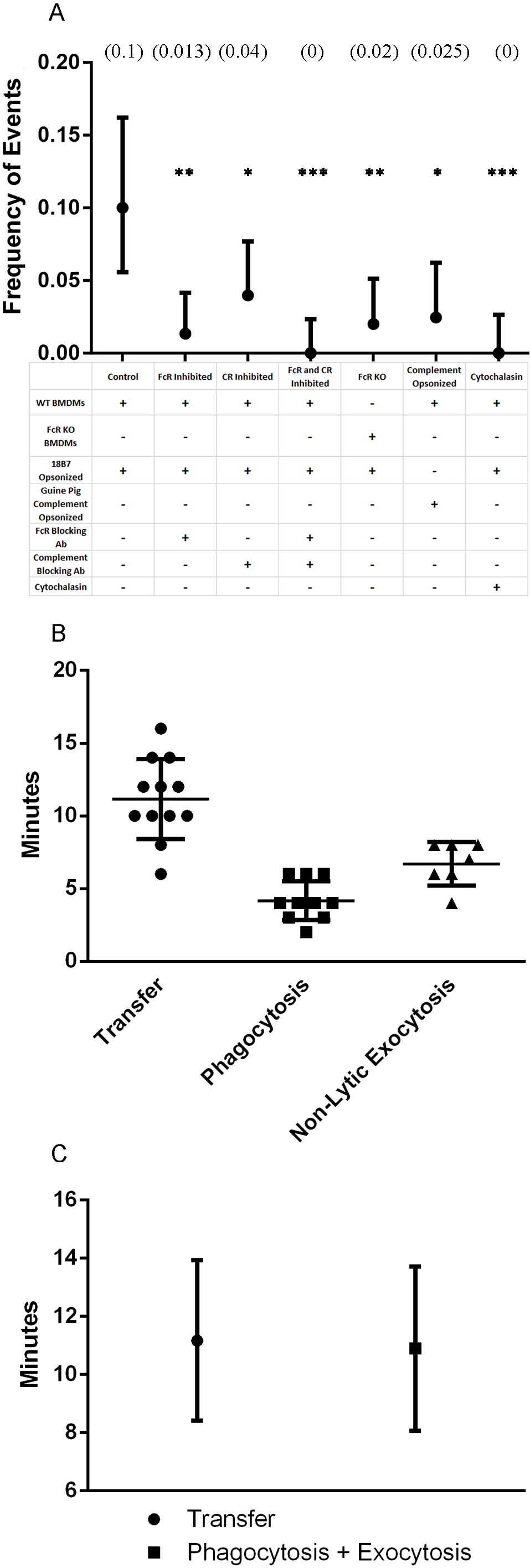
Inhibition and timing of transfer events supports exocytosis-phagocytosis hypothesis. **A.** Quantification of the frequency of transfer events observed between BMDMs supplemented with different inhibitor antibody combinations. Frequency values represent frequency of events based on number of events in all infected cells, with complete information found in Table I. **B.** Quantification of the total timespan of each transfer event as well as phagocytosis ingestion time determined by total frames captured. **C.** Quantification of the total timespan of transfer events compared to the addition of phagocytosis and exocytosis events. Data points represent population average with bars representing population standard deviations. P-value calculated as non-significant via unpaired two-tailed t-test.

**Table I.** Quantified cellular events in the presence or absence of antibody receptors. Counts are written as “Total # (Percent)”

| Event | Repeat Control | aCD16/32 | aCD11b | aCD16/32 + aCD11b | FcR KO | Complement Opsonized | Cytochalasin |
| --- | --- | --- | --- | --- | --- | --- | --- |
| **Lytic Exocytosis** | 12 (6.9) | 1 (0.67) | 0 | 1 (0.8) | 6 (4.03) | 9 (7.38) | 7 (6.25) |
| **Non-Lytic Exocytosis** | 9 (5.1) | 1 (0.67) | 4 (2.65) | 1 (0.8) | 4 (2.68) | 6 (4.92) | 0 |
| **Transfer** | 14 (8) | 2 (1.34) | 6 (3.97) | 0 | 3 (2.01) | 3 (2.50) | 0 |
| **Total Infected Cells** | 140 | 149 | 151 | 126 | 149 | 122 | 112 |

We hypothesized that the initial phagocytic event may have downstream effects on whether Cryptococcal cells can undergo cell-to-cell transfer. To explore a potential effect of complement mediated phagocytosis, we repeated these experiments with guinea pig complement opsonized *C. neoformans* on wild type BMDMs. We found that BMDMs which had ingested *C. neoformans* via complement experienced abrogated cell-to-cell transfer compared to control and in frequencies similar to both wild-type BMDMs inhibited with anti-FcR antibodies and Fcer1g BMDMs with no inhibiting antibodies (Figure 3A, Table I).

Finally, we also performed experiments supplemented with cytochalasin, an actin inhibitor, reasoning that actin activity is required for exocytosis. We observed no transfer events, with p < 0.001 compared to control, further supporting the idea that transfer relies on exocytosis events (Figure 3A). Given these results, we favored the first hypothesis as the continued presence of a phagosome around the cryptococcal cell would block antibody-receptor interactions.

### Temporal Kinetics of Cell-to-Cell Transfer

If Cryptococcal cells are transferred via exocytosis followed by phagocytosis it would follow that the total time of transfer events should resemble the length of exocytosis plus the length of phagocytosis. Cell-to-cell transfer time was estimated by counting the total number of frames immediately prior to immediately after transfer. Each image was taken at two-minute intervals so the total time of transfer was directly calculated by the number of frames. We found that the median and mean transfer times were 11 and 11.17 minutes, respectively, from twelve total analyzed events (Figure 3C). To our knowledge the timing of neither phagocytosis nor non-lytic exocytosis of *C. neoformans* by BMDMs has been previously carefully measured. To investigate whether the timing of cell-to-cell transfer matched the times of exocytosis and phagocytosis, we visualized phagocytosis by repeating the infection movie protocol but only adding opsonizing antibody immediately before starting image acquisition. We observed phagocytic events and counted the number of frames (taken each minute) to determine an experimental estimate of phagocytosis ingestion time. Based on eleven observed phagocytic events we determined ingestion occurred over approximately 4.18 minutes (Figure 3C). We then timed non-lytic exocytosis events. Based on seven observed non-lytic exocytosis events we determined a total expulsion time of approximately 6.71 minutes (Figure 3C). Adding the time required for non-lytic exocytosis (6.71 minutes) to that required for phagocytosis (4.18 minutes) yielded 10.9 minutes, which is tantalizingly close to the measured average time of 11.17 minutes for cell-to-cell transfer, p > 0.05 (Figure 3D).

### A Model for Macrophage-to-Macrophage Cellular Transfer

Our results indicate that cell-to-cell transfer is a coordinated process between two macrophages, that it does not involve the transfer of cytosol (e.g. Trogocytosis), and that it does not occur when the opsonic Fc and complement receptors are blocked. The time required for cell-to-cell transfer closely approximates addition of the times required for non-lytic exocytosis and ingestion. Although no single observation provides a mechanism, when our results are considered in combination, the most parsimonious interpretation is that *C. neoformans* cell-to-cell transfer results from sequential non-lytic exocytosis events followed by subsequent phagocytosis of expulsed yeast cells by an adjacent macrophage. (Figure 4).

**Figure 4.**
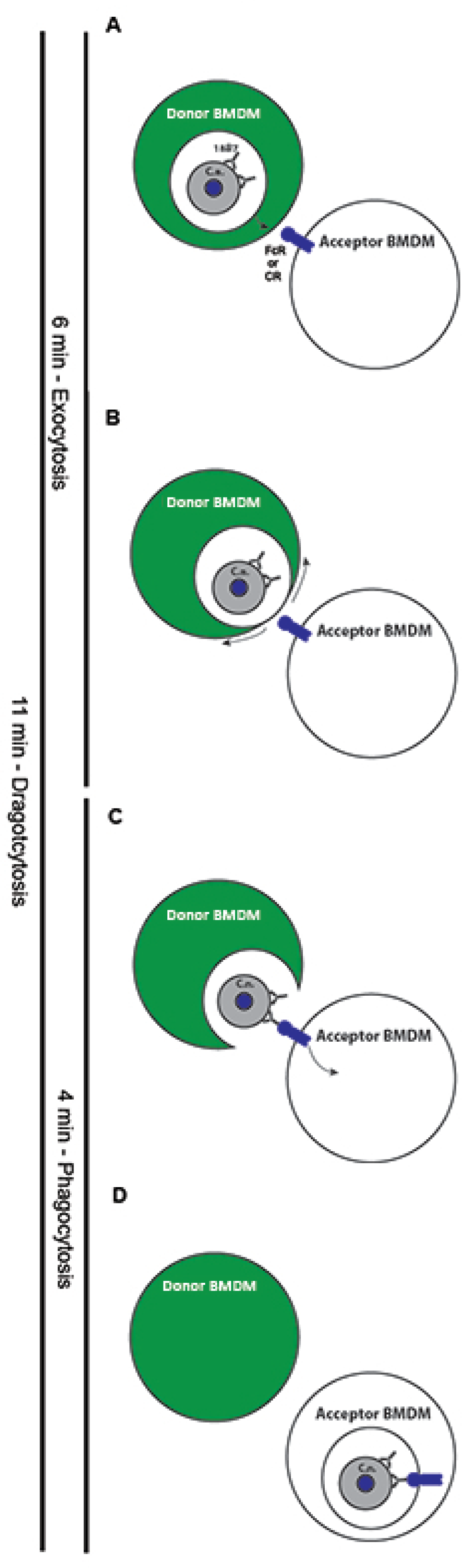
Proposed overarching model of cryptococcal cell-to-cell transfer. **A.** The initial step in transfer, and the point which was defined as the start of exocytosis. The moment when a phagosome (white space) containing an opsonized cryptococcal cell begins to move toward the plasma membrane of the donor macrophage. **B.** The second step of transfer when the phagosome reaches and merges with the plasma membrane of the donor macrophage in a manner which expels the cryptococcal cell. Cytosol is retained within the donor macrophage and excluded from the transfer process. The moment at which the cryptococcal cell is fully expelled from a macrophage was defined as the end of exocytosis. **C.** The third step of transfer in which the released cryptococcal cell is free to interact with the Fc receptor of the acceptor macrophage via opsonin which survives the initial phagosome. The moment of initial attachment of cryptococcal cell to macrophage was how the start of phagocytosis was defined. **D.** The final step of transfer, the point at which the cryptococcal cell has been fully ingested by the acceptor macrophage. This moment is also used to define the end of phagocytosis. **Below**. The total time of exocytosis, phagocytosis, and Dragotcytosis denoted by black line and measured time. The total length of Dragotcytosis roughly equals that of exocytosis plus phagocytosis.

## Discussion

Non-lytic exocytosis was implicated in persistence of fungemia(10) and may be involved in brain dissemination(23). Recently, non-lytic exocytosis was implicated as part of a process whereby phagocytic cells transfer *C. neoformans* to endothelial cells resulting in a new mechanism for cryptococcal crossing of the blood-brain barrier(16). Cell-to-cell transfer is a form of non-lytic exocytosis that has received relatively little attention, largely because it is difficult to study. In fact, the mechanism by which a fungal cell is transferred between two intact macrophages has been difficult to envision given that it involves a yeast cell in a membrane-bound phagosome that must cross two cellular membranes to travel from the donor to the acceptor cell.

We considered several hypotheses for the mechanism of cryptococcal cell-to-cell transfer. Initially we were intrigued by the possibility that cryptococcal cell-to-cell transfer occurred via Trogocytosis, a recently described cellular communication process in which cytosol and surface proteins are shared between macrophages(24). Trogocytosis was previously shown to be a mechanism that intracellular pathogens can utilize and promote to spread between macrophage cells(25). However, when we labeled the cytosol of macrophages and loaded them with cryptococcal cells we did not detect cytosol transfer from donor to acceptor cell during cryptococcal cell transfer. Shifts in the population data are clearly visualized in the accompanying density histograms which show no shift in cytosolic dyes before and after transfer events. It should also be noted that the shift in Uvitex 2B signal is immediately apparent despite the Uvitex 2B positive population (corresponding to the cryptococcal cell) accounts for a particularly small population of the entire region of interest (the entire macrophage). Therefore, even a small amount of cytosolic dye transferred to the donor cell would noticeably shift the population distribution post-transfer. Similarly, we could detect no transfer of plasma membrane signal from donor to acceptor macrophage in the experiments supplemented with membrane dye. The transfer experiments using cytosolic dye, combined with the lack of membrane transfer, essentially ruled out Trogocytosis as the mechanism of cryptococcal cell-to-cell transfer.

Consequently, we investigated whether cell-to-cell transfer was a non-lytic exocytosis event followed by ingestion of the expulsed yeast by a nearby macrophage. In support of this hypothesis was the observation that expulsed yeast cells have residual opsonizing antibody in their capsule even after phagosomal residence and that this antibody could support subsequent phagocytosis(21). When both FcR and CR were blocked no *C. neoformans* macrophage cell-to-cell transfer events were observed. Although our conditions should not have contained complement-derived opsonin, antibody binding to the *C. neoformans* capsule can result in structural changes that allow complement-independent phagocytosis through the complement receptor(22). These observations imply that cell-to-cell transfer is a sequential non-lytic exocytosis event followed by immediate phagocytosis from a nearby macrophage.

Non-lytic exocytosis events were reduced, but not completely abrogated, compared to controls when macrophages were incubated with FcR and CR blocking antibodies. This finding suggested that blocking antibody binding was somehow affecting exocytosis, raising the possibility that the reduced transfer in the presence of blocking antibody was a consequence of fewer donor cells. Consequently, we repeated these experiments with macrophages from Fc receptor knockout mice. As previously stated, opsonizing *C. neoformans* with 18B7 allows phagocytosis via the complement receptor so these macrophages can still be infected but will not be able to undergo cell-to-cell transfer relying on the Fc receptor. As hypothesized, we found that transfer events in Fc receptor knockout mice were reduced to levels similar to macrophages inhibited via antibody, while lytic and non-lytic events were not significantly lowered compared to control.

Based on the reduction but not complete inhibition of transfer events when only the Fc receptor was inhibited we decided to investigate the contribution of complement receptor, hypothesizing that it could be used as an alternative but less efficient cell-to-cell transfer route. Wild-type macrophages infected with guinea pig complement opsonized *C. neoformans* experienced cell-to-cell transfer rates similar to wild-type macrophages inhibited with anti-FcR antibody and macrophages from FcR knockout mice, while neither lytic nor non-lytic exocytosis events were significantly reduced compared to control. These data support our overall hypothesis that cell-to-cell transfer follows from coordination of a non-lytic exocytosis event followed by a phagocytosis event. Cell-to-cell transfer can be achieved through either Fc or complement receptor mediated phagocytosis, though Fc receptor mediated phagocytosis is more common.

An exocytosis-phagocytosis mechanism allows us to discard more complex explanations. For example, we had also hypothesized that the phagosome directly transferred between cells, but this also does not explain our results as inhibition of antibody receptors should not inhibit the direct transfer of an organelle. If the cryptococcal cells were retained in the phagosome throughout the cell-to-cell transfer process, then the phagosomal membrane would separate the opsonized cryptococcal capsule from Fc and complement receptors on the acceptor cell surface. Another hypothesis entertained was that cryptococcal cells are transferred utilizing membrane tunnel structures between macrophages. A tunnel transfer explanation is unlikely given that tunnels were absent from our microscopy analysis. In fact, when cell-to-cell tunnels occur these tend to be too small for even cytosolic molecules to diffuse through(26). In any case, a tunnel explanation is also ruled out by the fact that cell-to-cell transfer was abrogated by blocking cell surface opsonic receptors. Finally, due to the roles of Fc and complement receptors in immune synapse formation, we contemplated the possibility of cryptococcal cell transfer between macrophages via some type of previously uncharacterized macrophage specific immune synapse but concluded this would also be unlikely with respect to the data. Known immune synapses allow only small molecules and dyes to directly pass through. Particles larger than 32 nm are entirely excluded(27) which would also exclude cryptococcal cells.

The measured time of cell-to-cell transfer also supports the exocytosis-phagocytosis hypothesis. We determined the average length of non-lytic exocytosis events to be approximately 7 minutes, which is consisted with the prior reported time of 4-12 minutes(28). Macrophage phagocytosis of *C. neoformans* took approximately 4 minutes, providing measure for this process for the first time. Last, we measured that total cell-to-cell transfer took approximately 11 minutes, a time that appears close to the sum of macrophage phagocytosis and non-lytic exocytosis of *C. neoformans* times

We have termed the exocytosis-phagocytosis phenomenon Dragotcytosis, to denote the fact that this is a transfer between two sentinel (Greek ‘Dragot’) phagocytic cells. A new name would also allow us to distinguish it from other mechanisms denoting cell entrance or exit, such as: Trogocytosis, non-lytic exocytosis, and phagocytosis. Dragotcytosis differs from other mechanisms of cell to cell transfer mainly in that it involves the complete expulsion of cryptococcal cells from one macrophage before engulfment by another.

In summary, three lines of evidence indicate that cell-to-cell transfer represents sequential exocytosis-phagocytosis events: 1) we observed no cytoplasm transfer from donor to receptor cells; 2) the frequency of transfer events was greatly reduced by interference with receptors involved in phagocytosis; and 3) the time involved in cell-to-cell transfer was indistinguishable from the sum of the times involved in exocytosis and phagocytosis. The implications of this mechanism for pathogenesis and host defense are uncertain and may vary with the infection setting. Antibody administration can protect mice against cryptococcal infection(29). Comparative analysis of infected tissues of antibody-treated and control mice shows that the presence of antibody is associated with increased *C. neoformans* intracellular residence in macrophages(30). The increased prevalence of intracellular yeast cells was interpreted as reflecting more efficient antibody-mediated phagocytosis. However, our results suggest that Dragotcytosis could have contributed to this effect since fungal cells experiencing non-lytic exocytosis in granulomas could have been rapidly ingested by macrophages in close apposition to fungal infected cells. The fact that transfer events were completely absent between macrophages with inhibited antibody receptors has the interesting implication that these events may occur more frequently in hosts with robust antibody responses. Additionally, non-lytic exocytosis has been described with several other pathogens and Dragotcytosis could theoretically occur in other infections. For example, tuberculosis is a significant global disease in which intracellular pathogens can spread from lung granulomas, potentially through a similar mechanism. Whether Dragotcytosis benefits the host by promoting re-ingestion of expulsed cells or harms the host by promoting new cellular infections capable of spreading throughout the organism in a Trojan Horse manner is not known and probably depends on the circumstances when it occurs. Future research into Dragotcytosis should focus on elucidating what factors control the frequency or direction of transfer, whether there is a significant role for proximity of macrophages, whether there is a recruitment process for acceptor macrophages or specific surface markers upregulated by the donor, potential transcriptional differences between donor and acceptor macrophages or macrophages more likely to be donors vs acceptors, studying the process *in-vivo* for clinical relevance, and probing whether other relevant intracellular pathogens are capable of Dragotcytosis.

## Acknowledgements

Quigly Dragotakes would like to acknowledge Gundula Bosch Ph.D, Clive Shiff Ph.D., and David Sullivan M.D. for the timely assignment of a Trogocytosis paper in journal club during his rotation in the Casadevall Lab which inspired him to investigate this phenomenon.

Additionally, we would like to thank Carolina Coelho Ph.D. for sharing BMDM harvesting responsibilities on many occasions.

## Footnotes

1. The authors declare that we have no commercial or other association which might pose a conflict of interest.
2. Arturo Casadevall is supported by 5R01HL059842, 5R01AI033774, 5R37AI033142, and 5R01AI052733. Quigly Dragotakes is supported by a T32 and a fellowship from the ARCS-MWC foundation.
3. None of the data in this manuscript has been presented at any meetings.
4. All requests for reprints should be addressed to the corresponding author with information provided on the title page.
5. No authors affiliations have changed since completion.

**Supplemental Figure 1:** Compilation of density histograms representing the data portrayed in the cytosolic stain and Uvitex 2B quantifications. Vertical patterned bars represent the mean plus or minus the standard error of mean. These depictions offer a better visualization for the difference, or lack thereof, between stains before and after transfer events.

**Supplemental Figure 2:** Quantification of a plasma membrane marker before and after transfer events for donor and acceptor macrophages.

**Supplemental Movie 1:** A high resolution microscopy video of non-lytic cell-to-cell transfer of *C. neoformans* from one macrophage to another.

**Supplemental Movie 2:** A representative full move of one transfer event. Cells are visualized by phase contrast (gray), cytosolic stain (green), and Uvitex 2B (blue).

